# Establishing a Digital PCR-Based Reference Measurement Procedure for Monkeypox Virus: An Inter-laboratory Assessment and Standardization Study

**DOI:** 10.1101/2025.05.28.656494

**Authors:** Xia Wang, Huijie Li, Ruohui Guo, Yi Yang, Chunyan Niu, Shujun Zhou, Huafang Gao, Xiaohua Jin, Shangjun Wang, Meihong Du, Xiaoyan Cheng, Lingxiang Zhu, Lianhua Dong

## Abstract

The accurate detection of monkeypox virus (MPXV) is crucial for effective viral diagnosis and epidemic prevention. Currently, the field lacks standardized reference methods for MPXV quantification, as well as reliable pseudovirus reference materials (RMs) to ensure quality control throughout the detection process. To address this, we developed a droplet digital PCR (ddPCR) reference measurement procedure (RMP) for precise quantification of MPXV genes *B6R*, which demonstrates excellent performance with a wide dynamic range (11-1.24×10^4^ copies/μL), strong linearity (R^2^=0.9984), and a low limit of quantitation (11 copies/μL, CV≤25%). Repeatability tests confirmed high precision, with inter-laboratory CVs <10% across nine labs using different dPCR platforms. recovery efficiencies for *B6R* and *F3L* were ∼69%, with uncertainties incorporated into final measurements. The homogeneity assessment showed good results (CV=2.94%), and stability studies confirmed the stability of the sample during -70 °C and short-term storage (4 °C/-20 °C). Mandel’s statistical data shows that the reproducibility between laboratories is consistent. This validated ddPCR RMP provides metrological traceability, ensuring reliable and comparable results for MPXV diagnosis and supporting public health efforts.

## Introduction

Monkeypox virus (MPXV) is a zoonotic orthopoxvirus belonging to the *Orthopoxvirus* genus of the *Poxviridae* family. The virus was first discovered in laboratory monkeys in 1958 [1], MPXV was later detected in humans in 1970 in the Democratic Republic of Congo [2, 3]. Between 2022 and May 8, 2025, a total of 138,029 laboratory-confirmed cases and 317 deaths were reported worldwide. Structurally, MPXV is a double-stranded DNA virus with a brick-shaped or oval morphology, measuring approximately 200–250 nm in diameter. Its genome spans ∼197 kb and encodes roughly 190 proteins [4]. Similar to other orthopoxviruses, MPXV has high genetic stability, but can be divided into two main branches: the Central African (Congo Basin) branch and the West African branch, with the Central African branch being more pathogenic [5].

Transmission primarily occurs through zoonotic spillover, with limited human-to-human spread. Since May 2022, multiple countries have reported monkeypox cases in non endemic areas, indicating that the virus may have undergone epidemiological changes [6]. The incubation period after infection is usually 6-13 days (up to 21 days), with clinical manifestations including lymphadenopathy and a centrifugal rash progressing from the face to the extremities [7]. Monkeypox, as a serious zoonotic infectious disease, poses a certain threat to people’s health and public health safety. Its detection, diagnosis, and epidemic prevention and control are particularly important.

There are various methods for detecting monkeypox virus, and traditional methods include immunofluorescence using specific monoclonal antibodies [8], serum antibody ELISA using specific peptide segments [9]. These methods have problems such as cumbersome operation, long diagnostic time, high requirements for sampling environment, and easy sample contamination that affects experimental results. Molecular techniques, such as real-time PCR [10] and sequencing, offer alternatives, yet quantitative PCR (qPCR) remains constrained by amplification efficiency and reliance on external standards for relative quantification. In contrast, digital PCR technology (dPCR)enables absolute nucleic acid quantification, providing direct copy number measurements without calibration curves [11-14]. Despite these advances, the field lacks higher-order reference methods for MPXV quantification and pseudovirus reference materials (RMs) to ensure detection accuracy. In our previous study [15], a highly accurate dPCR method for MPXV *F3L* gene was established and was used for characterizing the reference material for *F3L* detection. However, the performance of the dPCR method in interlaboratory has not been evaluated.

In this study,, the aims of this study are to 1) establish a dPCR-based reference measurement procedure (RMP) for characterizing RMs for targeting the wild-type *B6R* of MPXV and 2) evaluate the inter-laboratory performance of the two proposed RMP for MPXV *F3L* and *B6R* by using the established reference materials.

## Materials and methods

### Preparation of Reference Materials (RMs)

This pseudo virus was synthesized by synthesizing the corresponding gene sequence of Monkeypox Virus *B6R* (Gene ID: 928902) in vitro, cloned and constructed into Ad5 replication defective adenovirus vector, and prepared as a pseudo virus in 293A cells [15]. The pseudo virus obtained after purification by chromatography column is a DNA sequence wrapped in adenovirus capsid, then diluted with virus protection solution to a certain concentration and divided into 110 μL per tube and stored at -70 °C.

### Establishment of digital PCR method

The primers and probes for *B6R* gene were designed with Primer Express 3.0.1 software (Table S1). Optimization of PCR reaction conditions for droplets digital PCR included annealing temperature, concentration of primer and probe. First, the final concentration of primers and probe under the condition of 400 nM and 300 nM, annealing temperature setting for 54 °C, 55 °C, 56 °C, 57 °C, 58 °C five annealing temperature. Then concentration of the primers and probe of *B6R*, concentration of the primer set up 300 nM, 400 nM, 500 nM, 600 nM, concentration of probe set up 200 nM, 300 nM, 400 nM, respectively. Data analysis was performed with the QuantaSoft software (version 1.7.4., Bio-Rad).

### Dynamic range, limit of quantification (LoQ), specificity and repeatability of dPCR

To verify the linear range of the established *B6R* method, DNA was diluted using a weight gradient method to obtain five concentrations. The linear range of the *B6R* gene was represented by (copies/μL). Perform 3 replicates for each dilution, and then calculate the consistency between theoretical and measured concentrations.

Analyze the coefficient of variation (CV) for each dilution and determine LoQ based on CV<25% [16]. To evaluate the method’s specificity, we employed the *B6R* and *F3L* methods to detect human cytomegalovirus (HCMV), human bocavirus (HBOV), as well as adenovirus types 7, 40, and 41(ADV7, ADV40 and ADV41),respectively. Repeatability refers to multiple consecutive repeated measurements of a target pseudovirus under the same measurement conditions. Repeat the experiment 3 times at three days, with at least 3 repetitions each time. The repeatability is represented by the CV of the measurement results.

### Recovery efficiency

The virus DNA was extracted by using the DNeasy Blood & Tissue Kit 250 (QIAGENC, #69506) according to the manufacture’s instruction. In order to evaluate the recovery efficiency of the recovery kit, the extracted DNA was used for secondary recovery. The established digital PCR calibration method was used to detect the concentrations before and after recovery, with 6 repeated recoverys. Each recovery was repeated three times, and the recovery efficiency was then calculated.

### Homogeneity and stability assessment

According to the number of units for packaging, 11 units were randomly selected from the initial, middle, and final stages of standard material packaging for homogeneity analysis. Perform three repetitions for each tube, and use *F*-test to calculate and analyze the results [17].

For short-term stability, three vials were analyzed at each testing time point (day 0, 3, 5, 7) under -20 °C and 4 °C, with a reference temperature of -70 °C. Long-term stability was measured at -70 °C for months 0, 1, 2, 3, 4, 8, 9, 21 and 33. Linear regression analysis was performed using time (*x*) as the horizontal coordinate and gene copy number concentration (*y*) as the vertical coordinate. It was evaluated according to ISO guide 35.

### Inter-laboratory assessment and characterization of the RMs

ISO 5725-1 [18] defines inter laboratory reproducibility as the accuracy of obtaining test results using the same method for the same test item in different laboratories and with different operators using different equipment. To maintain consistency, three vials of *B6R* RMs and the previous *F3L* RMs [15] were sent to nine laboratories and the absolute copy number of the samples was measured using the same detection method. Five laboratories conducted measurements on the BioRad platform (QX200), two laboratories conducted tests on Sniper DQ24, and Qiagen 24-well and SCI Digital each have one laboratory on the dPCR platform. Dixon’s and Grubbs’ tests are used to test for the presence of any outliers. According to ISO [19], Mandel’s *h* and *k* statistics were used to evaluate consistency between and within laboratories.

Uncertainty assessment of the *B6R* RMs includes the evaluation of uncertainties such as repeatability, homogeneity, and long-term storage stability. For the uncertainty of representation, the uncertainty of method accuracy, droplet volume [20], and recovery efficiency were considered. Calculate the extended uncertainty (*U*) using a coverage factor *k*=2.

## Results and Discussion

### DPCR optimization

As shown in the amplification result of Figure 1. According to the one-dimensional scatter plot of amplification, good amplification was obtained at all five annealing temperatures. Considering the repeatability of the results, 57 °C is determined as the optimal annealing temperature.

**Figure. 1.**
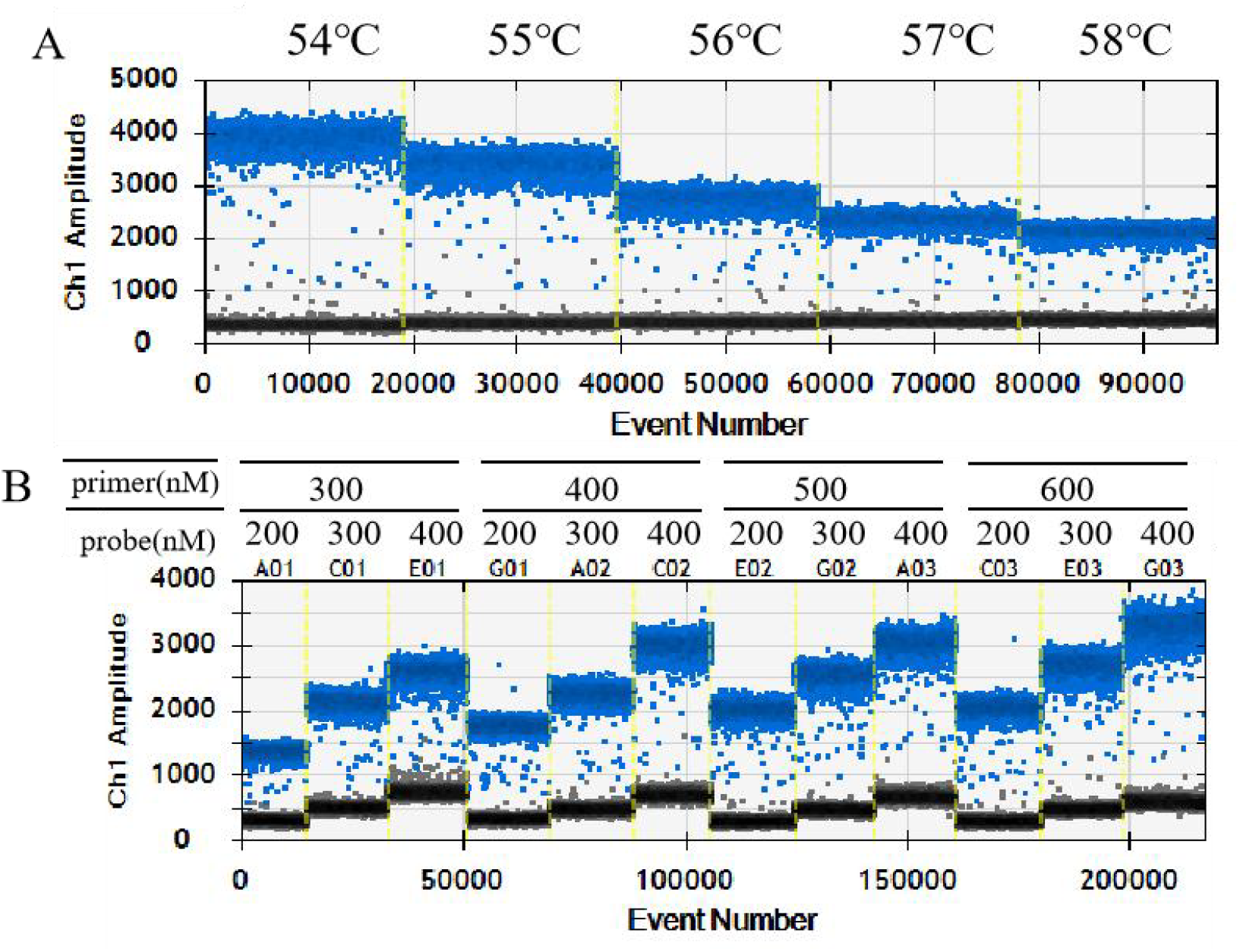
The results of dPCR optimization. A. Optimize the annealing temperature for the *B6R* assay. B. Optimize the concentration of primers and probes for the *B6R* assay.

When the primer concentration remains constant, the probe concentration of *B6R* increases from 300 nM to 600 nM, and the fluorescence intensity of both negative and positive droplets increases, as well as the degree of separation between negative and positive droplets. When the probe is at a constant concentration, the primer increases from 200 nM to 400 nM, and the fluorescence intensity of negative droplets remains basically unchanged. The luminescence intensity of positive droplets increases, resulting in an increase in the degree of separation between negative and positive droplets. However, as the probe concentration increases, the dispersion range of positive droplets becomes larger. For example, the distribution width of positive droplets at 300 nM is significantly larger than that at 200 nM. Based on the repeatability of quantitative results and copy number concentration factors, the primer concentration for *B6R* was determined to be 500nM, and the probe concentration was 300nM.

The total volume of the reaction system is 20 µL, including 10 µL of 2 × ddPCR Super Mix for Probe, primer and probe mixture, and 2 µL of DNA with a concentration of approximately 2500 copies/μL. PCR was performed at 95 °C for 10 minutes; 30 seconds at 94 °C, 1 minute at 57 °C, 40 cycles; Heat at 98 °C for 10 minutes using a thermal cycler (Veriti, Applied Biosystems).

### Dynamic range, limit of quantification (LoQ), specificity and repeatability of dPCR

The quantitative results of serial dilution measurements are shown in Table S2. As demonstrated in Figure 2, the measured target concentrations closely match the expected input values for each dilution. The ddPCR measurements show excellent linear correlation with theoretical DNA input values across five orders of magnitude (R^2^ = 0.9984, slope ≈ 0.9987), with exhibiting a linear dynamic range between 11 and 1.2 × 10^4^ copies/μL. Based on the coefficient of variation (CV) values shown in Table S2, a reaction containing an average of 11 copies/μL meets the limit of quantification (LoQ) criterion of CV ≤ 25%.

**Figure. 2.**
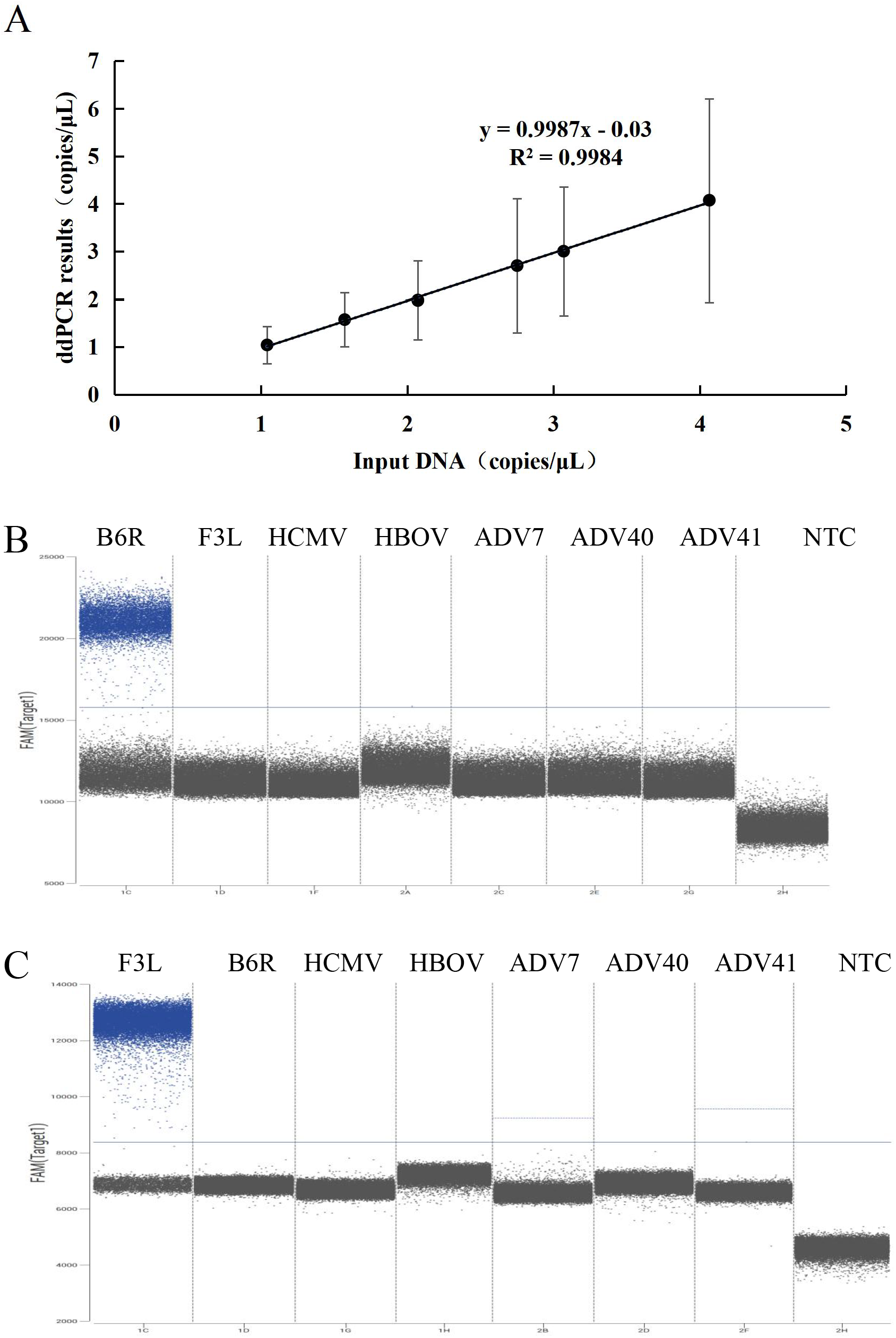
The results of the digital PCR method. A, the dynamic range of dPCR of *B6R* gene, with the coordinate axis based on the logarithm at the base of 10. B, the specificity of *B6R*; C, the specificity of *F3L*.

As shown in Figure 2B and 2C, As shown in Figure 2B and 2C, samples specifically amplified by the *B6R* and *F3L* assays showed no cross-reactivity with non-target viruses (HCMV, HBOV, ADV7, ADV40, and ADV41), confirming the high specificity of these methods. To define the repeatability, three measurements were performed on five samples in different runs on different days. The gravimetrically determined input values correlated well with dPCR-measured copy number concentrations for *B6R*, with an uncertainty ranging from 4.7% to 18% within the copy number concentration range of 11 to 1.2 × 10^4^ copies/μL (Table 1).

**Table 1.**
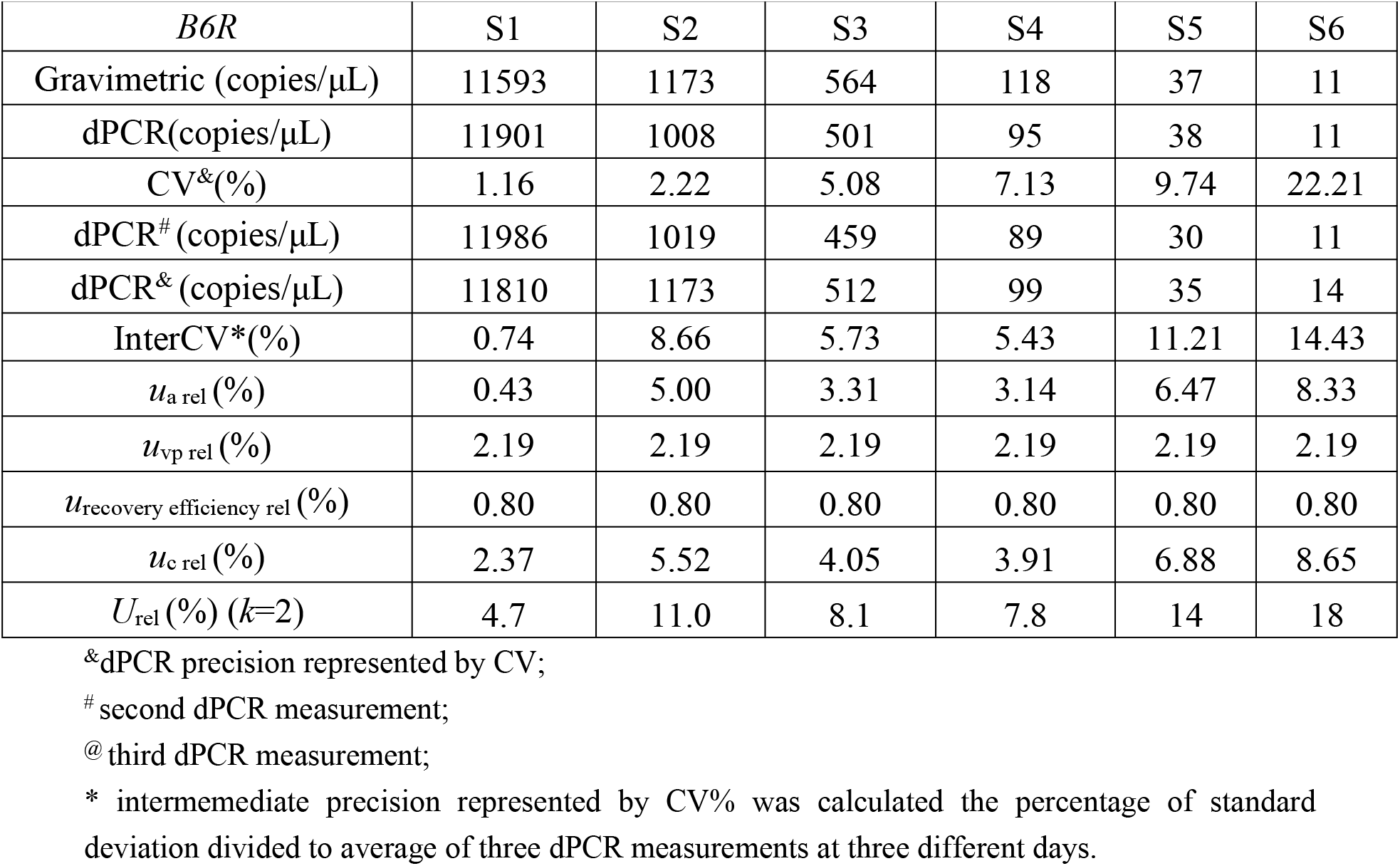
Measurement result and its uncertainty of *B6R* copy number concentration by dPCR.

### Recovery efficiency

In order to obtain the recovery efficiency of the reagent kit, *B6R* and *F3L* pseudo viruses were diluted to the detectable concentration range of digital PCR. The DNA extracted from *B6R* and *F3L* pseudo viruses was used as the original concentration, and the recovery efficiency was calculated by the ratio of the results of the secondary recovery to the original concentration. According to Table S3, the recovery efficiencies of *B6R* and *F3L* are 68.89% and 68.84%, respectively. The uncertainty associated with recovery efficiency was incorporated into the final measurement uncertainty.

### Homogeneity and stability assessment

Measured the homogeneity of the *B6R* RMs using dPCR and evaluate it according to ISO guide 35 [17]. The overall relative standard deviation between the vials was 2.94% (Figure 3). The *F*-test was used to calculate the homogeneity of *B6R* gene, and the results indicate that they are uniform because the obtained *F*-value is less than the *F*-critical value (Table S4). To ensure non-uniformity uncertainty, all relevant variables were considered, the uncertainty of non-uniformity was 1.18% (Table S4).

**Figure. 3.**
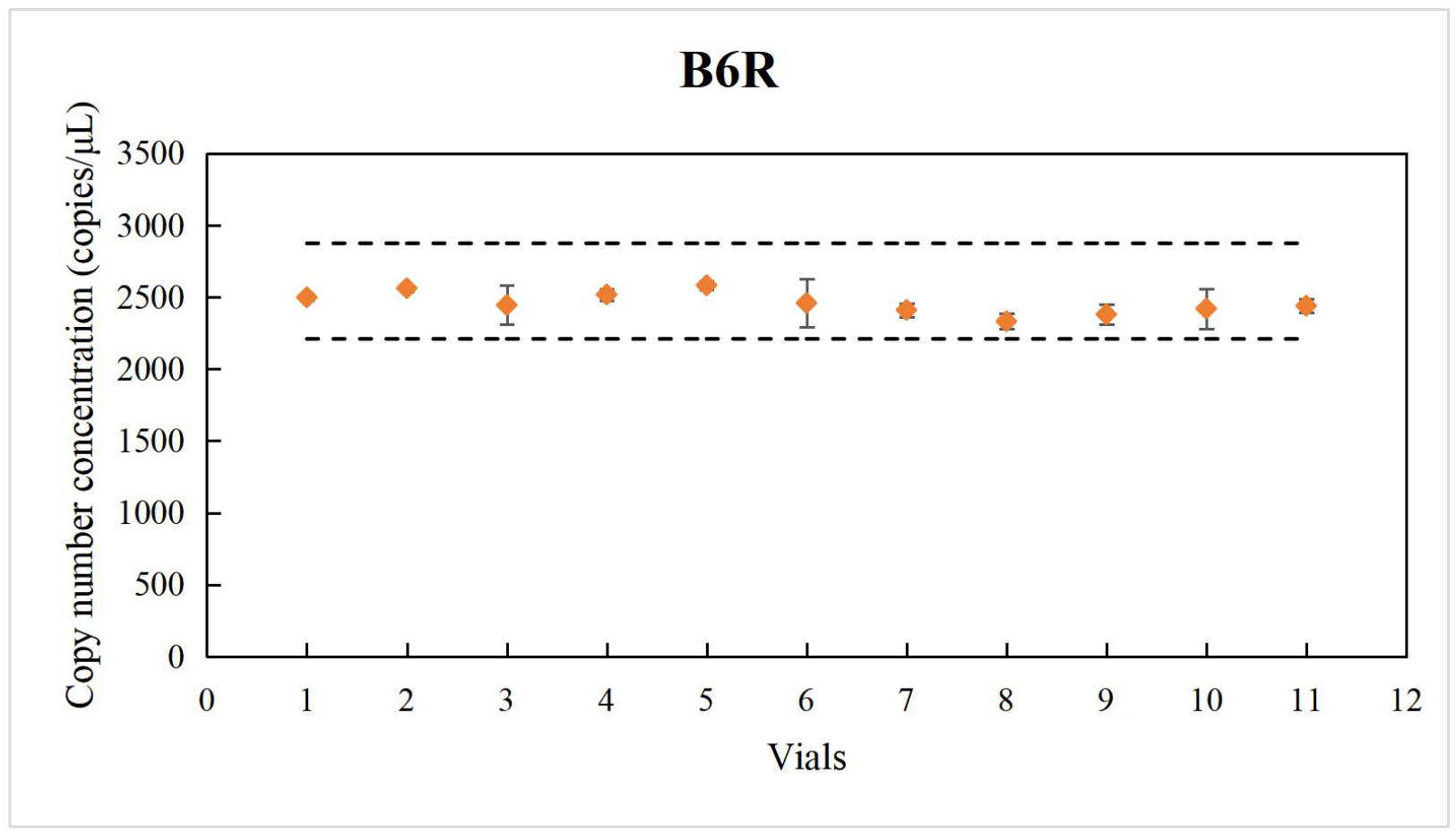
Homogeneity of the *B6R* RMs determined by dPCR assay.

The stability of *B6R* RMs was also measured by dPCR and evaluated according to ISO guideline 35. Long term stability analysis was conducted on the established method both *B6R* and *F3L* RMs. Linear regression analysis showed that no significant trend was observed at a 95% confidence level, indicating that the study sample remained stable at -70 °C. The relative uncertainty of long-term stability is shown in Table S5. For short-term stability of *B6R* RMs, no significant changes were observed within 7 days at 4 °C and -20 °C (Figure S1), indicating that the sample can remain stable at the test temperature during the one-week transportation period.

### Inter-laboratory assessment and characterization of the RMs

The inter laboratory results of 9 laboratories are consistent with the characteristic values within the extended uncertainty range (Figure S2). Five participating laboratories analyzed RMs using QX200, two using Sniper platform, and one using Qiagen and SCI Digital PCR platform. This QX200 and Sniper are based on droplet based digital PCR, while Qiagen and SCI Digital are based on chip based digital PCR. This enables further performance criteria for candidate reference measurement programs to be evaluated based on more dPCR platforms. The inter laboratory CV of *B6R* and *F3L* was within 10% (Table S6), and the reproducibility was calculated, indicating that the candidate PRMP has good reproducibility. The sources of uncertainty for the RMs mainly included the characterization methods, homogeneity and long-term stability of the *B6R* RMs (Table S7).

Calculate and plot Mandel’s *h* and *k* statistical data to evaluate inter laboratory and intra laboratory consistency, respectively (Figure 4). The Mandel diagram (Figure 4A) shows that all testing laboratories have consistent *B6R* copy number concentration measurements at the 1% significance level, indicating good inter laboratory reproducibility of *B6R*. Only laboratory 4 has significant variability at a significance level of 5%. The Mandel *k* plot of *B6R* reflects poor reproducibility observed at the 5% significance level for copy number concentration in laboratory 7, and consistency was achieved at the 5% significance level for reproducibility across nine laboratories (Figure 4B). For *F3L*, the consistency and repeatability of copy number concentration measurements at the 5% significance level were satisfactory across all laboratories, and no significant differences were observed between the 9 laboratories.

**Figure 4.**
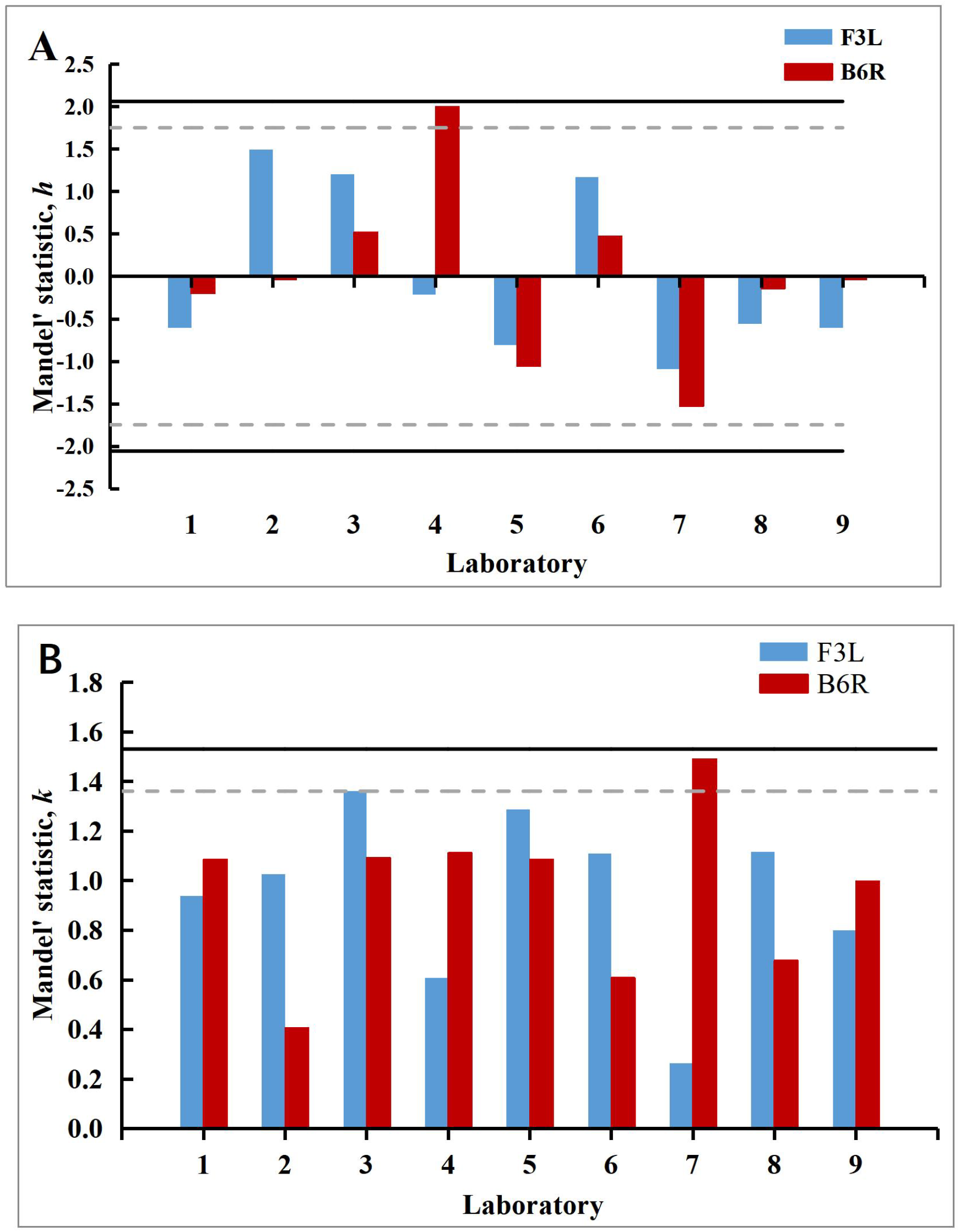
Consistency assessment by Mandel’s between-laboratory statistic, *h* (A) and within-laboratory statistic, *k*, (B), for the average number of measured copy concentration of *B6R, F3L* RMs, respectively. Lab 3,6: Sniper platform, Lab 4: SCI Digital platform, Lab 7: Qiagen 24-well platform, Lab 1,2,5,8,9: QX200 platform. The solid and dash lines are the 1% and 5% significance levels, respectively.

## Conclusion

The detection of monkeypox virus is particularly important for virus diagnosis and epidemic prevention and control, so it is necessary to develop accurate reference methods and RMs. Based on the optimal dPCR system, we established an RMP for quantifying *B6R* gene. The results indicate that our proposed dPCR RMP has high accuracy and good analytical sensitivity, and can be used as a reference method for monkeypox virus. Most importantly, it can be used to establish metrological traceability for monkeypox virus measurement, ensuring comparability and reliability of clinical diagnostic results.

## Acknowledgements

We appreciate the Turtle Technology Co., Ltd., QIAGEN Co., Ltd. and Sniper Medical Technology Co., Ltd. for their participation in the interlaboratory assessment.

## Funding

This work was supported by Basic scientific research project of National institute of metrology (AKYZD2202).

## Author contributions

Conceptualization and methodology, Lianhua Dong; Methodology, Lianhua Dong, Xia Wang, Huijie Li, Ruohui Guo, Yi Yang, Chunyan Niu, ; Investigation and data curation, Xia Wang, Huijie Li, Ruohui Guo, Yi Yang, Shujun Zhou, Huafang Gao, Xiaohua Jin, Shangjun Wang, Meihong Du, Xiaoyan Cheng, Lingxiang Zhu; Writing—original draft preparation, Lianhua Dong, Xia Wang; Supervision, Lianhua Dong. All authors have accepted responsibility for the entire content of this manuscript and approved its submission.

## Declarations

### Conflict of interest

The authors declare no competing interests.

## Notes

### Competing Interest Statement

The authors have declared no competing interest.

