## Supplementary materials for "Establishing a Digital PCR-Based Reference Measurement Procedure for Monkeypox Virus: An Inter-laboratory Assessment and Standardization Study"

### Table S1 Sequences and optimized concentrations of primer/probe of *B6R* gene

| Name | Primer/probe sequence | Final concentration |
| --- | --- | --- |
| *B6R*-F | CGATCTAACGAAGAATTTGATCCA | 500 nM |
| *B6R*-R | TCGAGAGTTTGCTCAGATCTGTCT | 500 nM |
| *B6R*-P | 5’-FAM- TGGATGATGGTCCCGAC -BHQ1-3’ | 300 nM |

### Table S2 The results of LoQ

| DdPCR results（copies/μL） | CV |
| --- | --- |
| 11901 | 1.16% |
| 1008 | 2.22% |
| 501 | 5.08% |
| 95 | 7.13% |
| 38 | 9.74% |
| 11 | 22.21% |
| 2 | 69.08% |

### Table S3 The results of recovery efficiency for *B6R* and *F3L*

|  | recovery efficiency | |
| --- | --- | --- |
| Times | *B6R* | *F3L* |
| 1 | 68.31% | 68.06% |
| 2 | 64.47% | 72.45% |
| 3 | 71.27% | 66.61% |
| 4 | 68.68% | 67.01% |
| 5 | 65.18% | 69.06% |
| 6 | 75.41% | 69.85% |
| Mean | 68.89% | 68.84% |
| CV | 5.36% | 2.84% |
| *u*_recovery efficiency ,rel_ | 2.19% | 1.16% |

### Table S4 The results of homogeneity for *B6R* RM

| Vial | repeat 1 | repeat 2 | repeat 3 |
| --- | --- | --- | --- |
| 1 | 2488 | 2474 | 2533 |
| 2 | 2591 | 2547 | 2547 |
| 3 | 2253 | 2518 | 2562 |
| 4 | 2488 | 2577 | 2488 |
| 5 | 2577 | 2547 | 2621 |
| 6 | 2695 | 2326 | 2356 |
| 7 | 2400 | 2474 | 2356 |
| 8 | 2326 | 2268 | 2400 |
| 9 | 2282 | 2444 | 2415 |
| 10 | 2385 | 2606 | 2268 |
| 11 | 2503 | 2415 | 2400 |
| Mean | 2458 | | |
| Q1 | 172833 | | |
| Q2 | 247026 | | |
| V1 | 10 | | |
| V2 | 22 | | |
| *F* | 1.54 | | |
| *F*_(10,22)_ | 2.30 | | |
| Conclusion | *F*<*F*_(10,22)_, reference material is homogeneous | | |
| *u_bb_*_,rel_ (%) | 1.18 | | |

### Table S5 The assessment of long term stability for *B6R* and *F3L* RM

| *B6R* | | *F3L* | |
| --- | --- | --- | --- |
| Time/month | dPCR (copies/μL) | Time/month | dPCR (copies/μL) |
| 0 | 2445 | 0 | 2907 |
| 1 | 2320 | 1 | 2934 |
| 2 | 2073 | 2 | 2711 |
| 3 | 2432 | 3 | 3026 |
| 4 | 2259 | 4 | 2768 |
| 8 | 2299 | 8 | 2857 |
| 9 | 2533 | 9 | 2763 |
| 21 | 2619 | 21 | 2823 |
| 33 | 2433 | 33 | 2577 |
| Mean | 2379 | Mean | 2818 |
| b_0_ | 2319 | b_0_ | 2890 |
| b_1_ | 6.74 | b_1_ | 7.93 |
| s | 154 | s | 107 |
| s(b_1_) | 4.92 | s(b_1_) | 3.41 |
| b_1_/s(b_1_) | 1.37 | b_1_/s(b_1_) | 2.32 |
| Freedom (n-2) | 7 | Freedom (n-2) | 7 |
| t_0.95,7_×s(b_1_) | 11.63 | t_0.95,7_×s(b_1_) | 8.07 |
| Result | \|b_1_\| < t_0.95, 7_×s(b_1_), stable | Result | \|b_1_\| < t_0.95, 7_×s(b_1_)，stable |
| *u*_s,rel_ (%) | 4.96 | *u*_s,rel_ (%) | 3.99 |

### Table S6 Interlaboratory result of *B6R* and *F3L* RM

| Lab | *B6R*（copies/μL） | *F3L*（copies/μL） |
| --- | --- | --- |
| 1 | 2490 | 3033 |
| 2 | 2532 | 3740 |
| 3 | 2675 | 3642 |
| 4 | 3053 | 3163 |
| 5 | 2271 | 2963 |
| 6 | 2663 | 3629 |
| 7 | 2150 | 2867 |
| 8 | 2505 | 3048 |
| 9 | 2532 | 3033 |
| **Mean** | **2541** | **3235** |
| SD | 241 | 317 |
| CV(%) | 9.5 | 9.8 |

### Table S7 Factors contributing to the relative standard uncertainty for MPXV reference material.

| **uncertainty** | ***B6R*** |
| --- | --- |
| *u*_a_^1^ (%) | 3.36 |
| *u*_v_^2^ (%) | 0.80 |
| *u*_f_^3^ (%) | 0.20 |
| *u*_recovery efficiency ,rel_ (%) | 2.19 |
| *u*_char_^a^ (%) | 4.09 |
| *u_bb_*^b^ (%) | 1.18 |
| *u_S_* ^c^ (%) | 4.96 |
| *u_c_*^d^ (%) | 6.54 |
| *U* (*k*=2)(%) | 13 |

^1^repeatability;^2^droplet volume;^3^dilution factor;^a^method characterization; ^b^homogeneity; ^c^stability; ^d^relative combined standard uncertainty; ^e^relative expanded uncertainty.

short-term stability

concentration(copies/μL)

| Time | -20℃ | 4℃ | 1 | 2 |
| --- | --- | --- | --- | --- |
| 0d | 2541.24 | 2541.24 | 2873.63 | 2208.85 |
| 3d | 2611.11 | 2613.43 | 2873.63 | 2208.85 |
| 5d | 2548.61 | 2513.89 | 2873.63 | 2208.85 |
| 7d | 2555.56 | 2480 | 2873.63 | 2208.85 |

### Figure. S1 Short-term stability of *B6R* RM

### Figure. S2 Interlaboratory assessment of candidate reference measurement procedure of *B6R* (Mean value± 2SD;the dashed line represents the range of uncertainty).

B6R

Copy number concentration(copies/μL)

| Laboratory |  | 1+多家定值图!$F$3:$F$13 |
| --- | --- | --- |
| 1 | 2490.28 | 2873.63 |
| 2 | 2532.22 | 2873.63 |
| 3 | 2674.82 | 2873.63 |
| 4 | 3052.76 | 2873.63 |
| 5 | 2270.93 | 2873.63 |
| 6 | 2662.83 | 2873.63 |
| 7 | 2150.29 | 2873.63 |
| 8 | 2504.8 | 2873.63 |
| 9 | 2532.22 | 2873.63 |
|  |  | 2873.63 |
|  |  | 2873.63 |

| Laboratory | 2 |
| --- | --- |
|  | 2208.85 |
| 1 | 2208.85 |
| 2 | 2208.85 |
| 3 | 2208.85 |
| 4 | 2208.85 |
| 5 | 2208.85 |
| 6 | 2208.85 |
| 7 | 2208.85 |
| 8 | 2208.85 |
| 9 | 2208.85 |
|  | 2208.85 |
